# Correlated patterns of genetic diversity and differentiation across an avian family

**DOI:** 10.1101/097733

**Authors:** Benjamin M. Van Doren, Leonardo Campagna, Barbara Helm, Juan Carlos Illera, Irby J. Lovette, Miriam Liedvogel

**Author notes:** Current address: Edward Grey Institute, Department of Zoology, University of Oxford, Tinbergen Building, South Parks Road, Oxford, OX1 3PS, UK.

## Abstract

Comparative studies of genomic differentiation among independent lineages can provide insights into aspects of the speciation process, such as the relative importance of selection and drift in shaping genomic landscapes, the role of genomic regions of high differentiation, and the prevalence of convergent molecular evolution. We investigated patterns of genetic diversity and divergence in stonechats (genus *Saxicola*), a widely distributed avian species complex with phenotypic variation in plumage, morphology, and migratory behavior, to ask whether similar genomic regions are important in the evolution of independent, but closely related, taxa. We used whole-genome pooled sequencing of 262 individuals from 5 taxa and found that patterns of genetic diversity and divergence are highly similar among different stonechat taxa. We then asked if these patterns remain correlated at deeper evolutionary scales and found that homologous genomic regions have become differentiated in stonechats and the closely related *Ficedula* flycatchers. Such correlation across a range of evolutionary divergence and among phylogenetically independent comparisons suggests that similar processes may be driving the differentiation of these independently evolving lineages, which in turn may be the result of intrinsic properties of particular genomic regions (e.g., areas of low recombination). Consequently, studies employing genome scans to search for areas important in reproductive isolation should account for corresponding regions of differentiation, as these regions may not necessarily represent speciation islands or facilitate local adaptation.

## INTRODUCTION

Evolutionary biologists seek to understand the genetic basis of speciation and the degree to which the divergence of lineages may involve independent changes on similar loci (Seehausen et al. 2014). Genomic sequencing has made it possible to examine patterns of differentiation across the genomes of organisms at different stages of divergence. Recent comparative studies of genome-wide patterns of variation, or “genomic landscapes,” have identified areas of the genome that are conspicuously differentiated relative to the genomic baseline among closely related taxa (Ellegren et al. 2012; Ruegg et al. 2014; Burri et al. 2015; Wang et al. 2016). It remains an open question whether these regions are functionally important in speciation, and whether they typically arise during speciation-with-gene-flow or as a consequence of selection in allopatry.

Most empirical studies have used F-statistics (Wright 1965) and other measures that compare allele frequencies between two populations to infer the magnitude of differentiation across the genome. These statistics are influenced by levels of within-population genetic variation and are therefore classified as “relative” measures of divergence (Hedrick 2005). Genomic outlier regions of high relative differentiation were first described as “islands of speciation” in the face of gene flow (Turner et al. 2005) and hypothesized to harbor loci that were important for reproductive isolation (Nosil et al. 2009; Feder et al. 2012). However, subsequent studies have identified alternate mechanisms by which isolated genomic regions of high differentiation can be generated in allopatry, and thus independently of gene flow (e.g., Noor and Bennett 2009; Turner and Hahn 2010; White et al. 2010; Cruickshank and Hahn 2014). For example, post-speciation selective sweeps or background selection, especially when combined with low recombination, can drive the differentiation of these loci relative to the rest of the genome (Nachman and Payseur 2012; Cruickshank and Hahn 2014; Burri et al. 2015).

Selective sweeps bring beneficial alleles to high frequency in a population, greatly reducing genetic diversity at linked sites via “hitchhiking” (Smith and Haigh 1974; Kaplan et al. 1989). The magnitude of the hitchhiking effect is influenced by recombination rate, with areas of low recombination experiencing greater linkage and a commensurate reduction in diversity across larger sections of a chromosome (Begun and Aquadro 1993; Charlesworth et al. 1997; Nielsen 2005). This reduction in local within-population genetic diversity results in high differentiation as measured by F_ST_ (Charlesworth 1998; Keinan and Reich 2010; Cruickshank and Hahn 2014). Recurrent sweeps in similar genomic regions across independent populations may be caused by selection for different advantageous alleles at the same locus. Alternatively, they could result from globally advantageous mutations transmitted among populations by gene flow, followed by sweeps due to local adaptation (see Roesti et al. 2014; Delmore et al. 2015). These processes can result in corresponding areas of low genetic diversity and high differentiation in independent population comparisons.

Similarly, background (or purifying) selection purges deleterious alleles as they arise and may also independently generate similar genomic landscapes of diversity and differentiation across populations (Charlesworth 2013; Burri et al. 2015; Wang et al. 2016). Under this scenario, a neutral variant that emerges in a population will persist only if it remains unlinked to a deleterious mutation. The removal of neutral variants linked to deleterious ones thereby reduces nucleotide diversity (Charlesworth et al. 1993; Charlesworth et al. 1997; Stephan et al. 1998). As recombination rates decrease, linkage extends over larger genetic distances and the probability of a neutral variant associating with a deleterious mutation (and thus being purged) is higher. Therefore, areas of low recombination will generally exhibit a greater reduction in genetic diversity due to background selection (Charlesworth et al. 1993; Nordborg et al. 1996). If highly conserved areas (e.g., of great functional importance) and/or areas of low recombination are similar across species, the effect of background selection may cause, or contribute to, parallel genomic landscapes of diversity and differentiation in comparisons of independently evolving lineages (Nordborg et al. 1996; Andolfatto 2001; Cruickshank and Hahn 2014).

The effects of background selection and selective sweeps on linked neutral loci have collectively been referred to as “linked selection” (Turner and Hahn 2010; Cutter and Payseur 2013; Cruickshank and Hahn 2014). The frequency with which parallel signatures of linked selection occur in closely related taxa and the contributions of background selection and selective sweeps in shaping genomic landscapes remain actively debated (Keinan and Reich 2010; Burri et al. 2015). In addition, the degree to which this parallelism may extend beyond a few well-studied species complexes is currently unknown.

Genome-wide scans of two independent groups of closely related bird species, *Ficedula* flycatchers and *Phylloscopus* warblers, have identified conspicuous peaks of relative divergence (i.e., genomic regions with very different allele frequencies) in pairwise comparisons of congeners that coincide with “valleys” of absolute divergence (i.e., regions with few sequence differences, not influenced by within-population genetic diversity) (Burri et al. 2015; Irwin et al. 2016). This inverse relationship is inconsistent with the speciation-with-gene-flow paradigm, where regions of high relative divergence are resistant to gene flow and therefore should show *high* absolute divergence (Noor and Bennett 2009; Nachman and Payseur 2012; Cruickshank and Hahn 2014). This suggests that post-speciation selection—not divergence-with-gene-flow—generates differentiation peaks in these systems. Within their respective species complexes, flycatchers and warblers show signatures of selection in similar genomic areas (Burri et al. 2015; Irwin et al. 2016), suggesting that in each group the same genomic regions may be important in multiple divergence events. However, neither the specific type of selection acting on these loci, nor their contribution to the speciation process, has been fully characterized. Furthermore, these studies primarily test the correspondence of divergent regions using correlation-based methods, which can be strongly affected by autocorrelation due to linkage.

Here, we investigate genome-wide patterns of molecular evolution in a well-studied group of birds to understand the extent to which similar genomic regions and similar evolutionary forces drive the divergence of closely related taxa. We focus on the stonechats (genus *Saxicola*), widespread insectivorous birds from open habitats of the Old World (Urquhart 2002; Collar 2016a, 2016b). Stonechats breed across a wide range of latitudes, from Southern Africa to Siberia, and thus experience a broad range of climatic conditions. Some taxa are restricted to small islands, while others span continents, and they range from long distance migrants to year-round residents (Baldwin et al. 2010). The genus *Saxicola* began diversifying during the late Miocene (8.2 million years ago; Illera et al. 2008) and currently comprises 15 recognized species (51 named taxa including subspecies; Gill and Donsker 2016). This well-documented evolutionary diversity makes *Saxicola* a powerful system for studying independently evolving lineages across a gradient of evolutionary differentiation, phenotypic variation, and life histories. Existing work on stonechats includes long-term studies of a range of phenotypic traits, from molt and metabolism to morphology and migration (Gwinner et al. 1983; Wikelski et al. 2003; Helm et al. 2005; Helm et al. 2009; Tieleman et al. 2009; Baldwin et al. 2010; Versteegh et al. 2014; Van Doren et al. In press). In addition, stonechats are phylogenetically close to *Ficedula*, a genus in the same family (Muscicapidae) that has been the focus of extensive research (Sætre and Sæther 2010). Here, we examine five stonechat taxa at disparate stages of divergence, including two that likely still exchange genes and two that diverged ~3.7 million years ago (Illera et al. 2008). These taxa show diverse morphological (e.g., wing and body size and shape) and behavioral (e.g., migratory direction and propensity) phenotypes. Furthermore, their demographic histories are likely varied: some reside on islands and others are continental.

We leverage the diversity of evolutionary histories in stonechats to investigate the extent to which genome evolution is correlated in independently evolving, but closely related, taxa. We ask: Have the same regions of the genome become differentiated over time in independent stonechat lineages? If so, what role has natural selection played in driving this correlated differentiation? We further ask if evolution is correlated at a deeper scale, between stonechats and two *Ficedula* flycatcher species. We posit that any loci that are differentiated in both genera are unlikely to arise from parallel ecological selection pressures and instead stem from intrinsic properties of those genomic regions. Finally, we examine the effects of life history and demography on the genome by comparing patterns of genetic diversity and differentiation between continental and island taxa. We hypothesize that an island taxon will show a weaker overall effect of selection on the genome, reflecting the theoretical prediction of increased drift with a smaller effective population size. Our goal is to shed light on the processes that influence the most conspicuous features—the high “peaks” and low “valleys”—of stonechat genomic landscapes. The results underscore the degree to which broad patterns of genetic diversity and differentiation are correlated across evolutionary time.

## METHODS

### Study system and sampling

We included five stonechat taxa in this study: *Saxicola rubicola rubicola* from Austria (European stonechat); *S. rubicola hibernans* from Ireland (European stonechat); *S. torquatus axillaris* from Kenya (African stonechat); *S. maurus maurus* from Kazakhstan (Siberian stonechat); and *S. dacotiae dacotiae* from Fuerteventura Island, Spain (Canary Islands stonechat) (Gill and Donsker 2016). Illera et al. (2008) estimated that African stonechats diverged from the remaining four taxa about 3.7 mya, Siberian stonechats subsequently split from the remaining three about 2.5 mya, and Canary Islands stonechats diverged from European stonechats about 1.6 mya. Illera and colleagues could not distinguish Austrian and Irish stonechats using mitochondrial DNA. We expect that Canary Islands stonechats have diverged from other taxa without gene flow because this taxon occurs on an oceanic island, and the present-day ranges of Kenyan and Siberian stonechats lead us to expect no ongoing gene flow between these and the other taxa. Conversely, we expect that Austrian and Irish stonechats likely still exchange genes because of their close geographic proximity, lack of mitochondrial divergence, and evidence of breeding dispersal between the British Isles and continental Europe (Helm et al. 2006).

Most stonechats included in this study originated from a common-garden experiment that Eberhard Gwinner initiated in 1981 at the Max-Planck Institute in Andechs, Germany, except for Canary Islands stonechats, which were directly sampled in the wild. For the other species, most birds were taken into captivity as nestlings, except for Irish stonechats (~50% captured on winter territories). The remaining sampled individuals (Table S1) were offspring of these captive stonechats, hatched between 1988 and 2006. Despite the inclusion of captive birds, relatedness within the pools was low (Table S1). Detailed descriptions of breeding and raising conditions are published elsewhere (Gwinner et al. 1987; Helm 2003; Helm et al. 2009).

The inclusion of second-generation progeny in our study could potentially lower measured levels of genetic diversity relative to a comparable sample of wild individuals. However, we find average genetic diversity (π) to be highest in Siberian stonechats, the species for which we incorporated the most captive-bred birds; this suggests that any putative bias is small and potentially negligible for the purposes of this study.

### Draft reference genome

We assembled the genome of a male Siberian stonechat (*S. maurus*) collected in Kazakhstan (44.59º N, 76.609º E) and housed at the Burke Museum (UWBM# 46478). We generated one fragment library with insert sizes of 180 base pairs (bp) and two mate-pair libraries (insert sizes: 3 and 8 kilobases), and we sequenced each of them on one Illumina HiSeq 2500 lane (obtaining 101-bp paired-end reads). We assembled the draft reference genome using the ALLPATHS-LG algorithm (Gnerre et al. 2011) and used HaploMerger (Huang et al. 2012) to improve the assembly by merging homologous scaffolds and removing those resulting from the erroneous split of two haplotypes into separate scaffolds. The final Siberian stonechat assembly comprised 2,819 scaffolds, with a total scaffold length of 1.02 Gb and an N50 scaffold size of 10.0 Mb. Half of the final assembly is represented in 24 scaffolds, and 75% in 65 scaffolds. Ambiguous bases (N’s) make up 4.4% of its total length.

We assembled scaffolds from our stonechat reference genome into draft chromosomes by mapping them to the *Ficedula albicollis* genome assembly, version 1.5 (RefSeq accession GCF_000247815.1) (Kawakami et al. 2014) and used SatsumaSynteny (Grabherr et al. 2010) to align the *Saxicola* draft genome to the *F. albicollis* assembly. Because these species are phylogenetically close and synteny is relatively conserved among birds (Ellegren 2010), this method allowed us to position 97.1% of the stonechat reference genome in the presumed correct order. Inversions and other chromosomal rearrangements occur in birds (Backström et al. 2008), so it is possible that a small percentage of the genome may be ordered or oriented incorrectly.

### Resequencing of five stonechat taxa

We extracted genomic DNA from stonechat blood or tissue samples using a salt extraction protocol and selected 49-56 individuals (n_total_ = 262, including both males and females) from each of the five stonechat taxa for sequencing (Table S1). We created five pooled libraries, one per taxon, from equimolar aliquots of DNA using the Illumina TruSeq DNA kit and sequenced them on an Illumina NextSeq 500 (Table S2).

We used BWA-MEM (Li 2013) to align sequences to our reference genome and performed refinement and quality control steps with Picard (http://broadinstitute.github.io/picard/) and the Genome Analysis Toolkit (GATK) (McKenna et al. 2010), including filtering by mapping quality and removing duplicate sequences (Supporting information). Sequences mapped to the draft reference genome at a mean per-pool coverage between 13.8x and 26.1x, and mean mapping quality was between 45-46 for all taxa (Table S2). Although this level of coverage is insufficient to sequence every individual at every locus, the goal of our pooled sequencing strategy was to estimate population allele frequencies by sampling a subset of chromosomes in a pool. Gautier et al. (2013) found that allele frequencies of individual SNPs estimated with 10-50x pool coverage (pool size = 30) were strongly correlated with estimates derived from separate individual-based (n = 20) sequencing at 1x-6x (r = 0.93) and 6-10x (r = 0.94) per individual. Additionally, the effects of pool-derived sampling error are greatly reduced in window-based analyses where variation and differentiation are summarized across groups of SNPs (Kofler et al. 2011a). Because we use a windowed approach and therefore do not rely on the frequencies of individual SNPs, we are confident that we can accurately assess and compare genome-wide patterns of genetic variation with this level of coverage.

### SNP-based phylogeny and inter-taxa divergence

Although we used an existing mitochondrial phylogeny (e.g., Illera et al. 2008) as a basis for our study approach and design, we also constructed a phylogenetic tree for the five focal stonechat taxa using nuclear markers. This step was designed to confirm the mitochondrial findings and serve as a basis for phylogeny-based inference. We used Pied and Collared Flycatchers *(Ficedula hypoleuca* and *F. albicollis*) as outgroups. We constructed a maximum likelihood tree with RAxML (Stamatakis 2014), using 17,527,493 fixed single-nucleotide polymorphisms (SNPs) from across the nuclear genome (Supporting information). We applied the Lewis correction, following the recommendation of Stamatakis (2014), for ascertainment bias resulting from the exclusion of invariant sites.

We generated mean genome-wide F_ST_ and d_XY_ pairwise distance matrices for the five focal stonechat taxa and displayed them graphically using principal coordinate analyses performed with the *ape* package in R (Paradis et al. 2004).

### Pooled population genomic analyses

We analyzed sequence data with the software packages *npstat* (Ferretti et al. 2013) and *Popoolation2* (Kofler et al. 2011b), designed specifically for the analysis of pooled sequencing data. With *npstat* we calculated: (1) Tajima’s *D*, to test for rare variants as a signal of directional or balancing selection or large-scale demographic effects; (2) π, an estimate of genetic diversity, which is derived from the number of pairwise sequence differences among members of a population; and (3) Fay and Wu’s *H*, a statistic related to Tajima’s *D* but sensitive only to high frequency derived alleles, thus influenced by positive selection but not by background selection (Fay and Wu 2000). We polarized alleles using the Collared Flycatcher.

We then used *Popoolation2* to calculate pairwise F_ST_ among all pairs of taxa. We also estimated d_XY_ (Nei and Li 1979; Cruickshank and Hahn 2014), a measure of absolute divergence, as *A_X_B_Y_* + *A_Y_B_X_*, where *A* and *B* are the frequencies of the two alleles at a locus and *X* and *Y* denote the two groups being compared.

For all analyses, we excluded bases within 5 bp of indels to reduce the probability of including erroneous genotypes due to misalignments. We calculated all metrics for 50-kb non-overlapping windows (Supporting information), within which we only considered sites with minor allele counts ≥ 2 and coverage between half and three times that of the pool’s average. We only retained windows in which at least 40% of bases (i.e. 20 kb) satisfied this coverage criterion. For d_XY_, we calculated the windowed value by summing over the window and dividing by the total number of sites with sufficient coverage (variable or not).

### Correlation analyses

We first quantified the similarity of genome-wide patterns of genetic diversity and divergence using Spearman rank correlations. Although the p-values of these tests are affected by autocorrelation due to genetic linkage, they are nonetheless a valuable summary of genome-wide similarity and provide a means to compare the results of the present study with previous work.

### Identification of genomic outlier regions

We identified regions of the stonechat genome showing consistently elevated or lowered values of Tajima’s *D*, π, Fay and Wu’s *H*, F_ST_, and d_XY_, and therefore may be important in the divergence of stonechat lineages. In particular, we wanted to determine whether any genome-wide similarities revealed by the correlation analyses were driven by a relatively small number of genomic regions. We applied a kernel-based smoothing algorithm across 50-kb windows (box density with bandwidth of 30; see Supporting information) and compared this smoothed line with 25,000 smoothed lines obtained after permuting the order of the windows (see Ruegg et al. 2014). We called outlier locations where the observed smoothed line was more extreme than the most extreme smoothed value from the null (permutation) distribution. We merged outlier regions separated by four windows or fewer (i.e. by <200 kb). Because the effective population size of the Z chromosome is smaller than that of the autosomes, baseline levels of variation and differentiation are different from those of autosomes (Charlesworth 2001). We therefore permuted windows of the Z chromosome and autosomes separately (see Fig. S1, Supporting information).

### Concordance of genomic outlier regions within *Saxicola*

Once outlier regions were identified, we assessed their overlap among stonechat taxa. For each pairwise comparison, we counted the number of outlier regions that showed any degree of overlap between the two datasets. By considering each region separately, we account for the autocorrelation among their constituent windows.

We then tested whether the observed number of overlapping regions was significantly greater than expected under the null hypothesis of no association in outlier positions between datasets, using a custom permutation test. While holding the number and size of outlier regions constant, we randomly permuted their locations across the genome 1,000 times and measured the degree of overlap under these simulated scenarios. The p-value of the test was the proportion of simulations under which the number of overlapping outlier regions was equal to or greater than the observed value; we thus accounted for the varying number and size of outlier regions in each comparison. We applied a false discovery rate correction to each series of tests (Benjamini and Hochberg 1995).

For each comparison, we calculate the proportion of outlier regions in one genomic landscape also present in the other (and vice versa) and report the greater of these two values. Thus, if landscape 1 shows 10 peaks and landscape 2 shows 50 peaks, and 9 out of 10 peaks in landscape 1 are also present in landscape 2, our outlier similarity score will be 9/10 = 0.90.

Previous studies have used scatterplots and correlation analyses as the primary manner of assessing association between outlier regions in independent comparisons (e.g., Burri et al. 2015; Irwin et al. 2016). However, these tests are affected by autocorrelation due to genetic linkage. By considering each outlier region as a single unit, our permutation approach overcomes this issue by treating each contiguous outlier region (instead of each 50-kb window) as an independent observation.

### Correspondence of genomic landscapes between *Saxicola* and *Ficedula*

To test for conservation of genomic landscapes at a deeper level of divergence, we compared stonechat genomic landscapes to those of the genus *Ficedula*. We calculated F_ST_, d_XY_, and π for Pied Flycatchers (*F. hypoleuca*) and Collared Flycatchers (*F. albicollis*). We downloaded reads from the Sequence Read Archive (project ERP007074; accession PRJEB7359; http://www.ncbi.nlm.nih.gov/sra) for 10 individuals of each species (Smeds et al. 2015) (Table S3), and processed reads for quality as described for the stonechat analysis (Supporting information).

We filtered, trimmed and aligned *Ficedula* reads to the stonechat draft reference genome so that we could directly compare the locations of outlier regions between genera, using the same tools in GATK and Picard as for stonechat sequences (Supporting information). To calculate F_ST_, we first generated a VCF file with UnifiedGenotyper from GATK and filtered raw variants with the VariantFiltration tool (settings: QD < 2.0 || FS > 60.0 || MQ < 40.0). We then calculated F_ST_ with VCFtools (Danecek et al. 2011) from the resulting SNPs across 50-kb non-overlapping windows. We estimated d_XY_ from minor allele frequencies obtained in ANGSD (Korneliussen et al. 2014), using a custom script to calculate 50 kb windowed averages. We only included sites that had genotype calls for at least 5 out of 10 individuals per species and retained windows for which at least 40% of bases satisfied this criterion.

## RESULTS

### SNP-based phylogeny and inter-taxa divergence

The Maximum Likelihood (ML) phylogeny built on fixed nuclear sites showed high support for the placement of the Canary Islands stonechat as the sister taxon to the European stonechat (Austria and Ireland) (Fig. 1A). The clade comprising European, Canarian, and Kenyan stonechats, to the exclusion of Siberian stonechats, was also strongly supported. This nuclear phylogeny contradicted the existing mtDNA topology. We verified that this result was not an artifact of sparse taxon sampling or choice of outgroup by constructing an ML tree with cytochrome-*b* consensus sequences obtained from Austrian, Irish, Kenyan, and Siberian pools; not enough mitochondrial sequence was recoverable for Canarian stonechats. Here, Kenyan stonechats were placed as the sister lineage to the remaining stonechats, in agreement with past mitochondrial studies (not shown).

**Figure 1.**
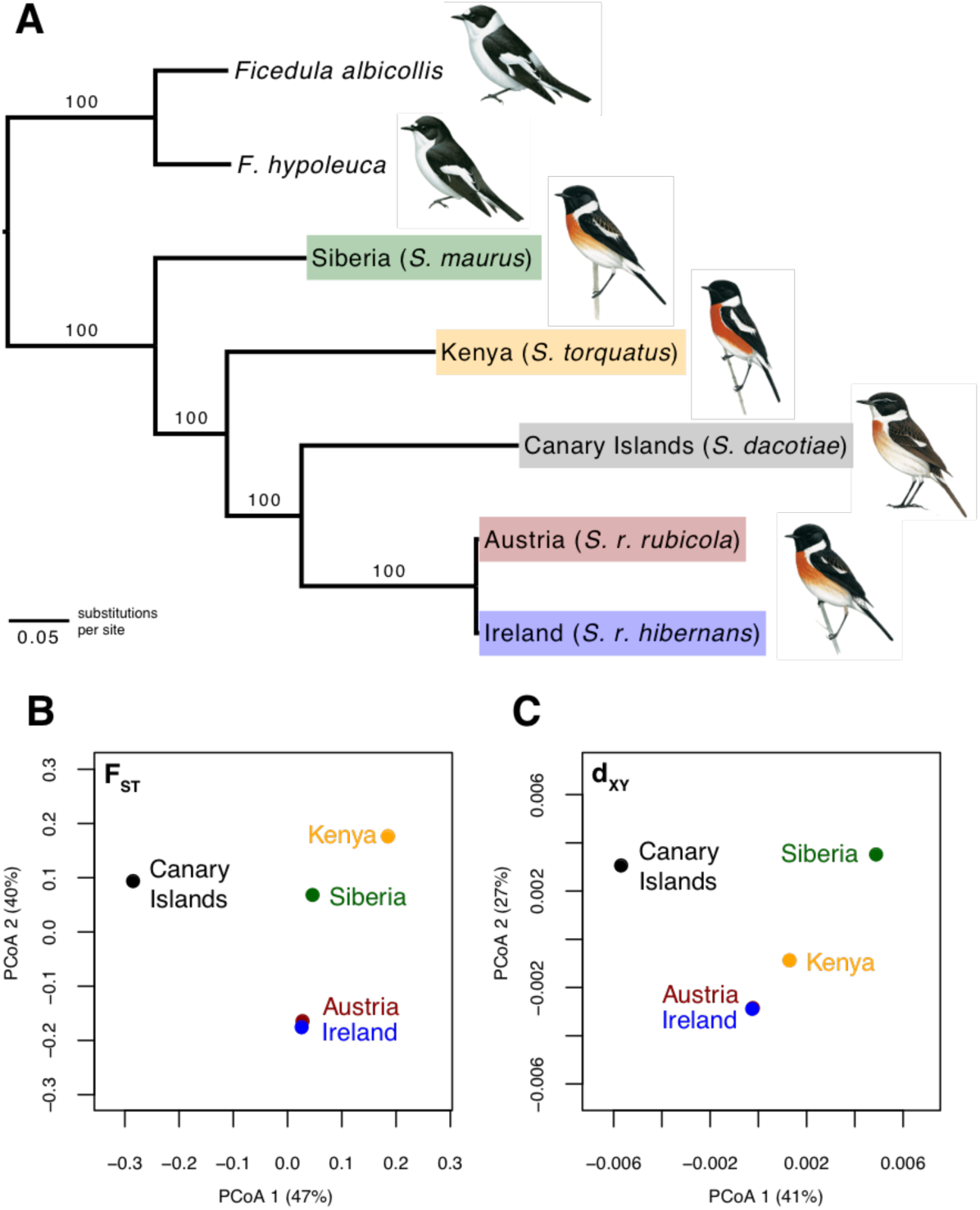
(A) Maximum likelihood phylogenetic tree constructed with RAxML from fixed sites across the stonechat nuclear genome, with two Ficedula species used as outgroups. Branch labels denote bootstrap support from 100 rapid bootstrap iterations. This topology places Siberian stonechat (S. maurus) as the sister lineage to the remaining taxa, in contrast to previous trees based on mitochondrial DNA. Canary Islands stonechats are most closely related to European stonechats. Illustrations (males shown) are reproduced with permission from Handbook of Birds of the World Alive (Collar 2016a, 2016b). (B, C) Biplot of two principle coordinate axes derived from analyses of: (B) mean F_ST_ and (C) mean d_XY_. Axes are labeled with percent of variance explained.

We calculated mean genome-wide F_ST_ and d_XY_ to further examine relationships among stonechat taxa. The first two principal coordinate axes calculated from a distance matrix of mean pairwise F_ST_ (explaining a total of 87% of variance; 47% in first axis) revealed Siberian stonechats to be approximately equidistant from the other taxa in terms of overall allele frequency differentiation (Fig. 1B). Stonechats from Austria and Ireland were extremely similar (with only 7 fixed differences out of 10,164,331 sites with F_ST_ > 0, or 7 × 10^−5^ %), reflecting their geographic proximity and common evolutionary history. In contrast, Kenyan and Canary Islands stonechats were most different (1,251,605 fixed differences out of 12,401,462 sites with F_ST_ > 0, or 10.09%). Overall, Canary Islands stonechats were strikingly dissimilar to even their closest evolutionary relatives (Austria vs. Canary Islands: 782,967 fixed differences from 12,754,086 variable sites, or 6.14%). European stonechats were more similar genome-wide to Siberian and Kenyan stonechats than to those from the Canary Islands, their sister lineage (Austria vs. Siberia: 244,623 fixed out of 15,168,199 variable, or 1.61%; Austria vs. Kenya: 640,425 fixed out of 12,032,148 variable, or 5.32%).

The principal coordinate analysis based on d_XY_ (68% of variance explained by first two axes; 41% by the first) was similar to the one based on F_ST_, except that Kenyan stonechats were closer to European stonechats than to Siberian stonechats (Fig. 1 C). This is consistent with the nuclear tree (Fig.1 A). Again, Austria and Irish stonechats were nearly identical. Canary Islands stonechats were distant from all stonechat taxa, but most similar to the European taxa.

### Shared regions of high differentiation show low genetic diversity, except in Canary Islands stonechats

Measures of divergence were strongly correlated among stonechats. D_XY_ showed strong correlations across genomic windows (Fig. 2 A-B), and d_XY_ outlier regions were highly similar (Fig. 2 D; Figs. S2 and S3, Supporting information); mean outlier similarity scores, averaged across all comparisons, were 0.85 for low d_XY_ outliers and 0.79 for high d_XY_ outliers (Fig. S4, Supporting information). F_ST_ was also significantly correlated in all comparisons, but the strength of this correlation varied (Fig. 3 A-D). The association was greatest in comparisons including Siberian stonechats (Fig. 3 A), but F_ST_ was also correlated among independent comparisons (i.e. with no shared taxon) (Fig. 3 B-C). Overall F_ST_ outlier similarity was lower than d_XY_ for both peaks and valleys (means of 0.31 and 0.24, respectively), indicating that approximately one-third of F_ST_ peaks were shared (Fig. 3 E and Fig. S5, Supporting information). Across all comparisons, windows with the lowest F_ST_ showed the most consistent associations. Of note, outlier regions showed significant overlap in several comparisons where the four taxa being compared were all different (Fig. 3 E), implicating common processes in independent stonechat lineages in the generation of differentiation landscapes.

**Figure 2.**
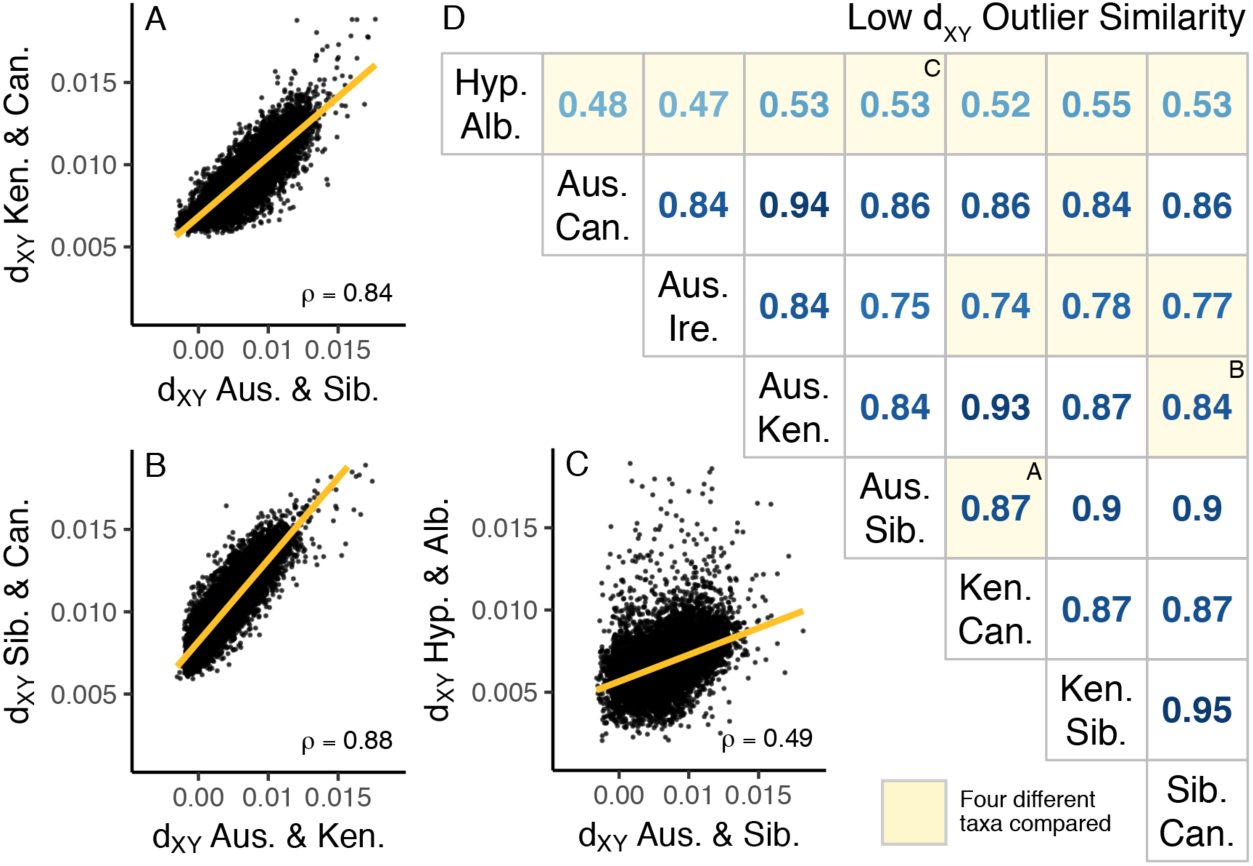
Correlation of d_XY_ among stonechats and flycatchers (Ficedula hypoleuca and albicollis, or “Hyp.” and “Alb.”). (A,B,C) show scatterplots where each point represents one 50-Kb genomic window. Orange lines are best-fit lines, and the Spearman’s rank correlation rho (ρ) coefficient is given. (D) shows outlier similarity scores, which quantify the number of low-d_XY_ “valleys” shared among different comparisons. Some comparisons including Irish stonechats are not shown because of their similarity to Austrian stonechats. All tests were significant after applying a false discovery rate correction. Cells with yellow backgrounds indicate that four independent taxa are being compared. Letters in the upper right of cells show which cells correspond to the scatterplots in sections A-C.

**Figure 3.**
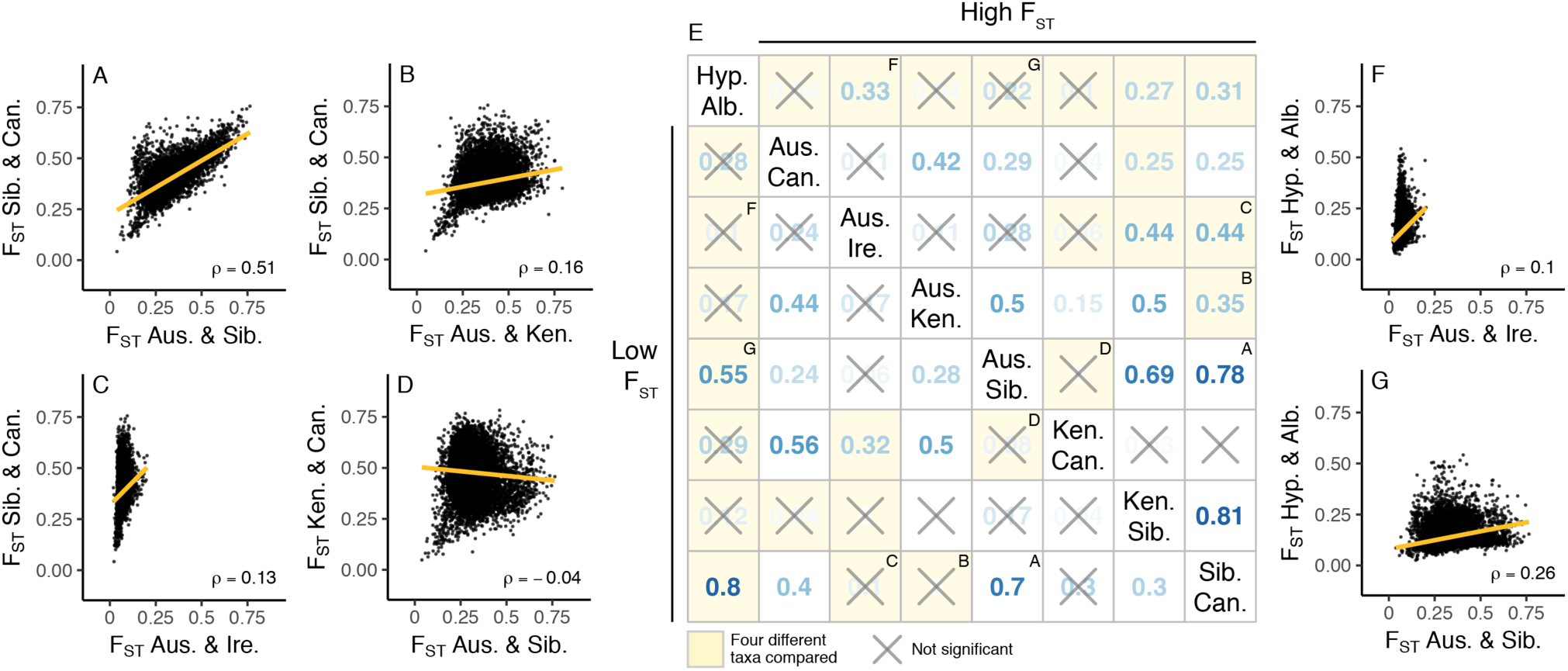
Correlation of F_ST_ among stonechats and flycatchers (Ficedula hypoleuca and albicollis, or “Hyp.” and “Alb.”). (A,B,C,D,F,G) show scatterplots where each point represents one 50-Kb genomic window. Orange lines are best-fit lines, and the Spearman’s rank correlation rho (ρ) coefficient is also given. (E) shows outlier similarity scores, which quantify the number of high-F_ST_ “peaks” shared among different comparisons (upper triangle of matrix) and the number of low-F_ST_ “valleys” shared among different comparisons (lower triangle of matrix). Some comparisons including Irish stonechats are not shown because of their similarity to Austrian stonechats. Cells with an ‘X’ indicate tests that were not significant after applying a false discovery rate correction. Cells with yellow backgrounds indicate that four independent taxa are being compared. Letters in the upper right of cells show which cells correspond to the scatterplots in the other sections.

Generally, regions of high F_ST_ showed low genetic diversity, both within (π) and between (d_XY_) stonechat taxa. F_ST_ and d_XY_ were strongly negatively correlated, especially in comparisons including Siberian stonechats (Fig. 4 A-B), and F_ST_ peaks overlapped strongly with d_XY_ valleys (Fig. 4 A). Regions of reduced absolute divergence also showed reduced nucleotide diversity (Fig. 5). Reductions in diversity occurred in the same genomic windows among stonechats, even after standardizing for levels of between-population diversity (Fig. 6; Figs. S6 and S7, Supporting information). Note that, due to the high similarity between Austrian and Irish stonechats, we do not present comparisons of Irish stonechats with non-Austrian stonechats.

**Figure 4.**
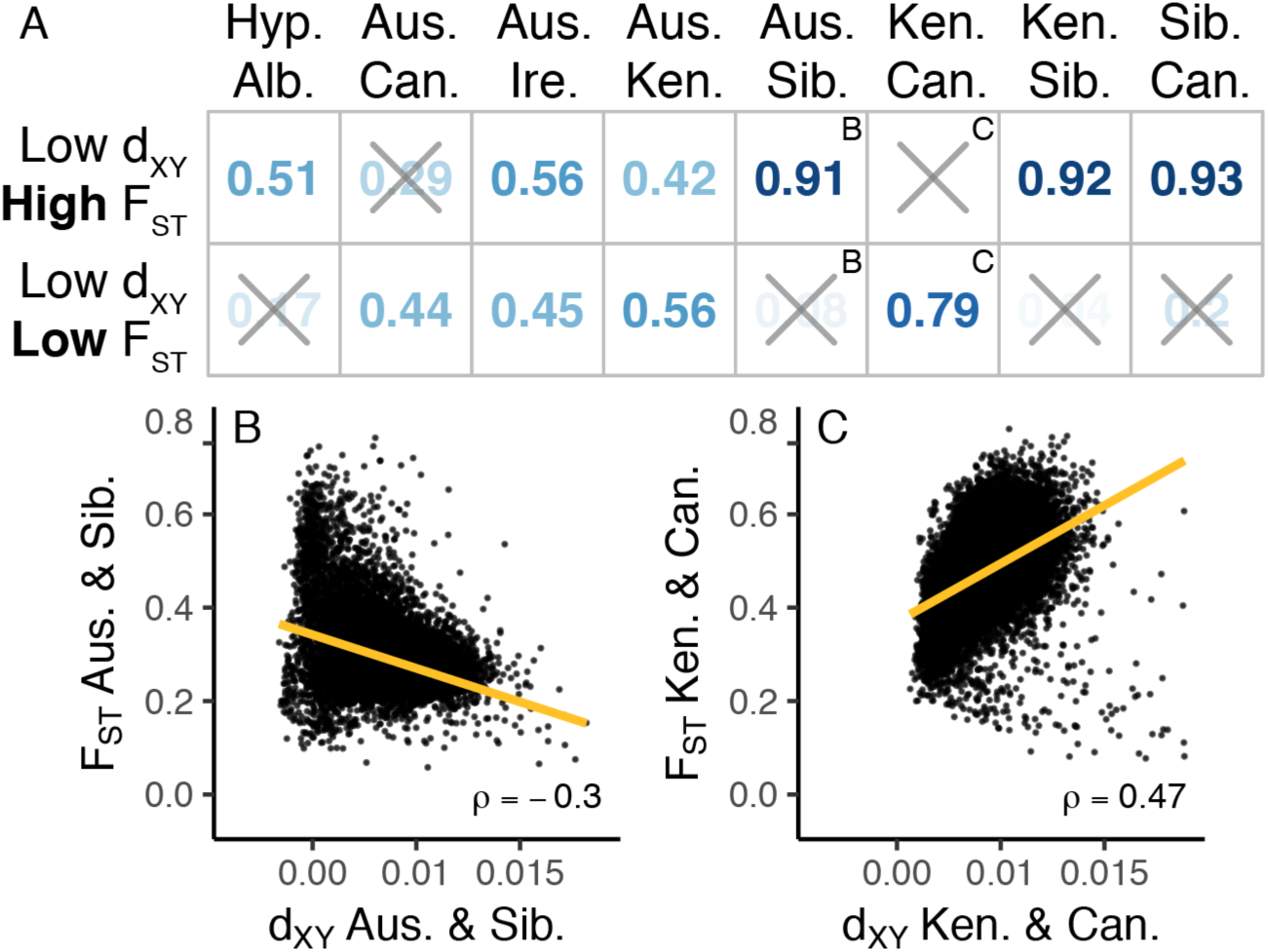
Correlation of F_ST_ and d_XY_ among stonechats and flycatchers (Ficedula hypoleuca and albicollis, or “Hyp.” and “Alb.”). (A) shows outlier similarity scores, which quantify the number of low-d_XY_ “valleys” that coincide with either high-F_ST_ “peaks” (top row) or low-F_ST_ “valleys” (bottom row). (B,C) show scatterplots where each point represents one 50-Kb genomic window. Refer to Figs. 2–3 for details.

**Figure 5.**
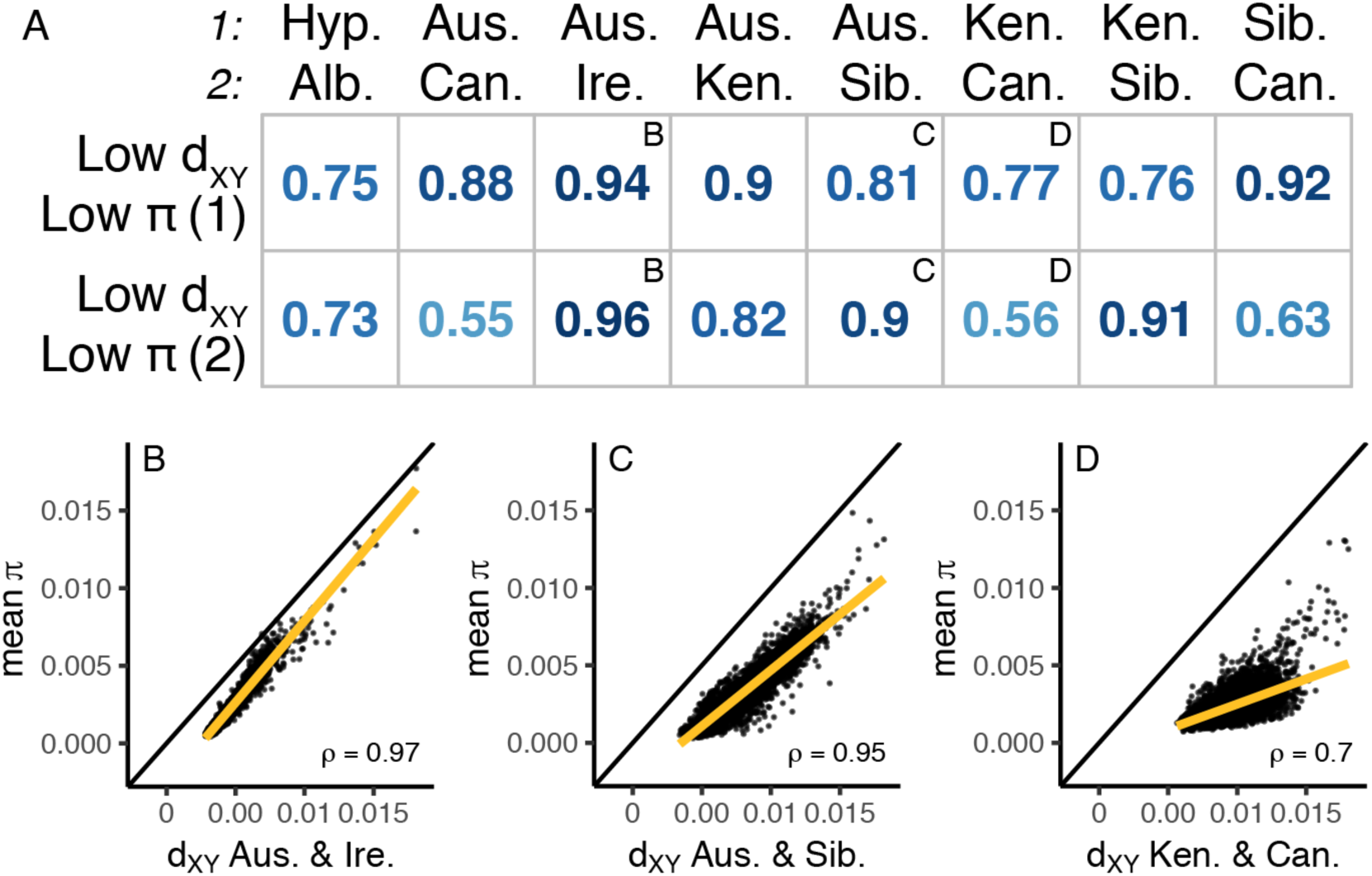
Correlation of nucleotide diversity (π) and d_XY_ among stonechats and flycatchers (Ficedula hypoleuca and albicollis, or “Hyp.” and “Alb.”). (A) shows outlier similarity scores, which quantify the number of low-d_XY_ “valleys” that coincide with low π “valleys” in the two taxa being compared (top row and bottom row). The top (No. 1) and bottom (No. 2) rows show the results for each of the two taxa being compared. This is necessary because π is a single-population statistic, while F_ST_ and d_XY_ compare two populations. All comparisons were significant after applying a false discovery rate correction. (B,C,D) show scatterplots where each point represents one 50-Kb genomic window. Refer to Figs. 2–3 for details.

**Figure 6.**
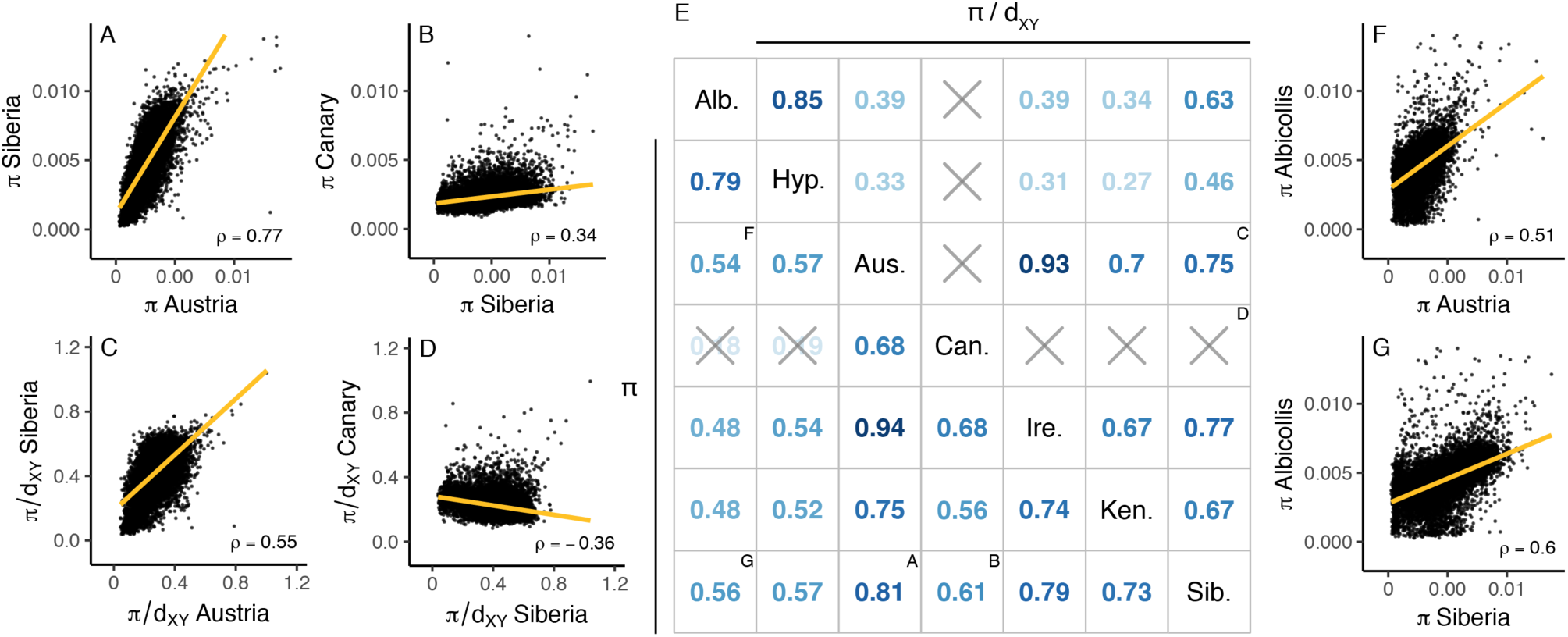
Correlation of nucleotide diversity (π) and standardized nucleotide diversity (π/d_XY_) among stonechats and flycatchers (Ficedula hypoleuca and albicollis, or “Hyp.” and “Alb.”). (A,B,C,D,F,G) show scatterplots where each point represents one 50-Kb genomic window. (E) shows outlier similarity scores, which quantify the number of π/d_XY_ valleys shared among different comparisons (upper triangle of matrix) and the number of π valleys shared among different comparisons (lower triangle of matrix). Refer to Figs. 2–3 for details.

However, these patterns of genetic variation were often weakened, absent, or even reversed for Canary Islands stonechats. Across the genome, F_ST_ and d_XY_ were *positively* correlated, despite the lowest-F_ST_ windows showing high d_XY_ (Fig. 4 C). Canary Islands stonechats showed the weakest associations in diversity correlations (Fig. 5 A,D, Fig. 6 B,D). Standardized nucleotide diversity (π/d_XY_) was negatively correlated between Canary Islands and Siberian stonechats (Fig. 6 D), indicating that the regions of the Siberian stonechat genome that showed the greatest diversity reductions were, in fact, relatively more diverse in Canary Islands stonechats than the rest of the genome (Fig. S7, Supporting information). No π/d_XY_ valleys regions were shared between Canarian stonechats and other stonechat taxa (Fig. 6 E).

### Evidence of selection and effects of demography

Among stonechats, genomic regions of high differentiation (F_ST_), low absolute divergence (d_XY_), and low genetic diversity (π) coincided with significant decreases in Tajima’s *D* and Fay and Wu’s *H* (Figs. 7 and 8). Fay and Wu’s *H* showed strong associations with F^ST^ only in comparisons including Siberian stonechats. Fay and Wu’s *H* outlier regions were relatively infrequent but coincided with low Tajima’s *D* and π when they occurred, except in Canary Island stonechats (Fig. S8, Supporting information). Tajima’s *D* outlier regions were generally shared across stonechats (Fig. S9, Supporting information), Some distinct low-H outlier regions occurred in only one taxon (e.g., chromosomes 4A and 6) (Fig. S10, Supporting information). Overall, in addition to lacking genetic diversity, outlier regions contained more low frequency alleles than the rest of the genome, which is highly suggestive of a role of natural selection in shaping differentiation patterns.

**Figure 7.**
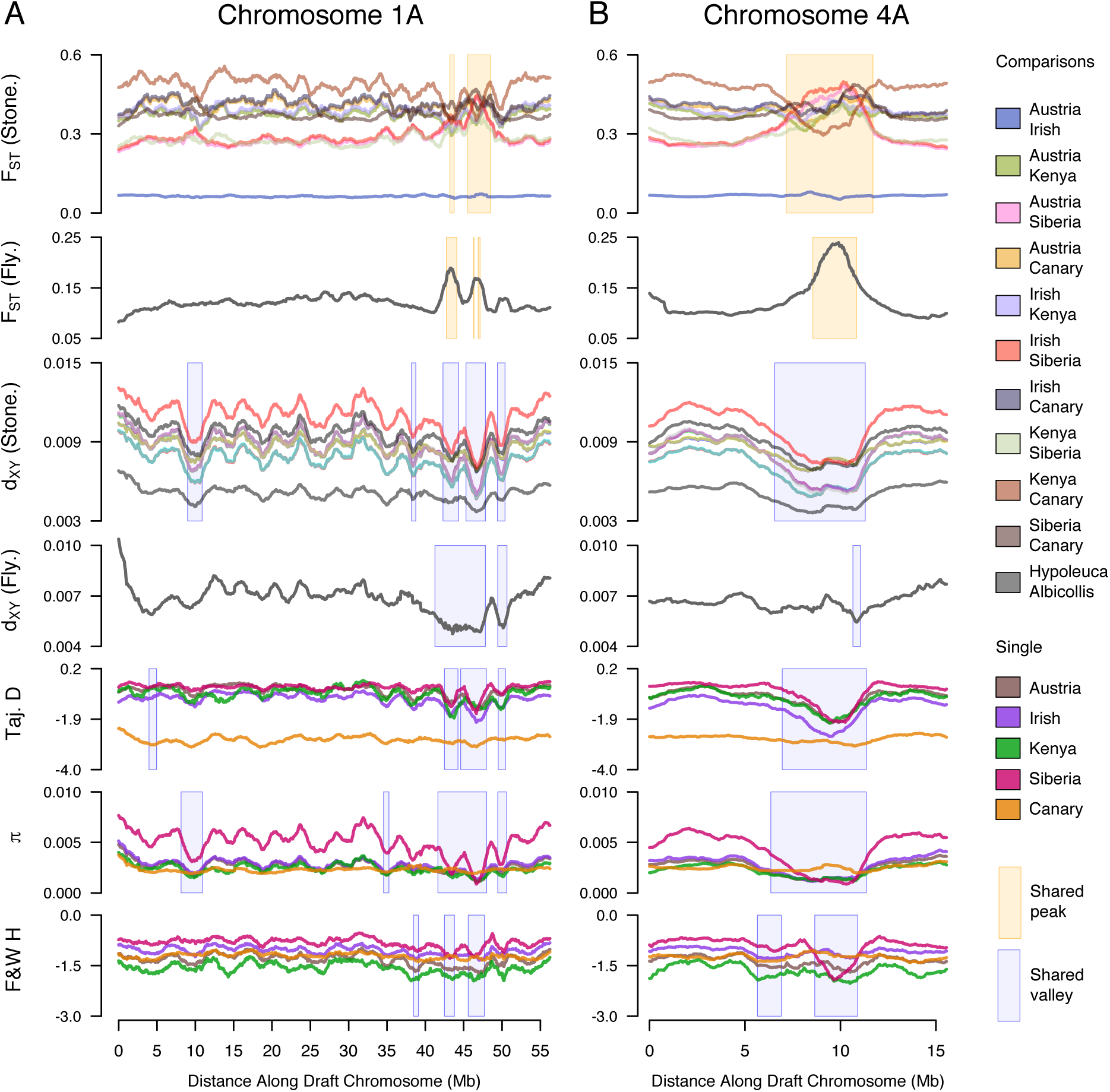
Genomic statistics calculated across stonechat and flycatcher chromosomes 1A and 4A. Yellow and blue boxes indicated shared peaks and valleys, respectively. From top to bottom, the statistics and box details are: F_ST_ among stonechats (peaks shared by 3 or more comparisons), F_ST_ between flycatchers (peaks that also overlap with shared stonechats peaks), d_XY_ among stonechats (valleys shared by 5 or more comparisons), d_XY_ among flycatchers (valleys that also overlap with shared stonechats valleys), Tajima’s D (valleys shared by 2 or more taxa), nucleotide diversity (π) (valleys shared by 2 or more taxa), and Fay & Wu’s H (valleys shared by 2 or more taxa).

**Figure 8.**
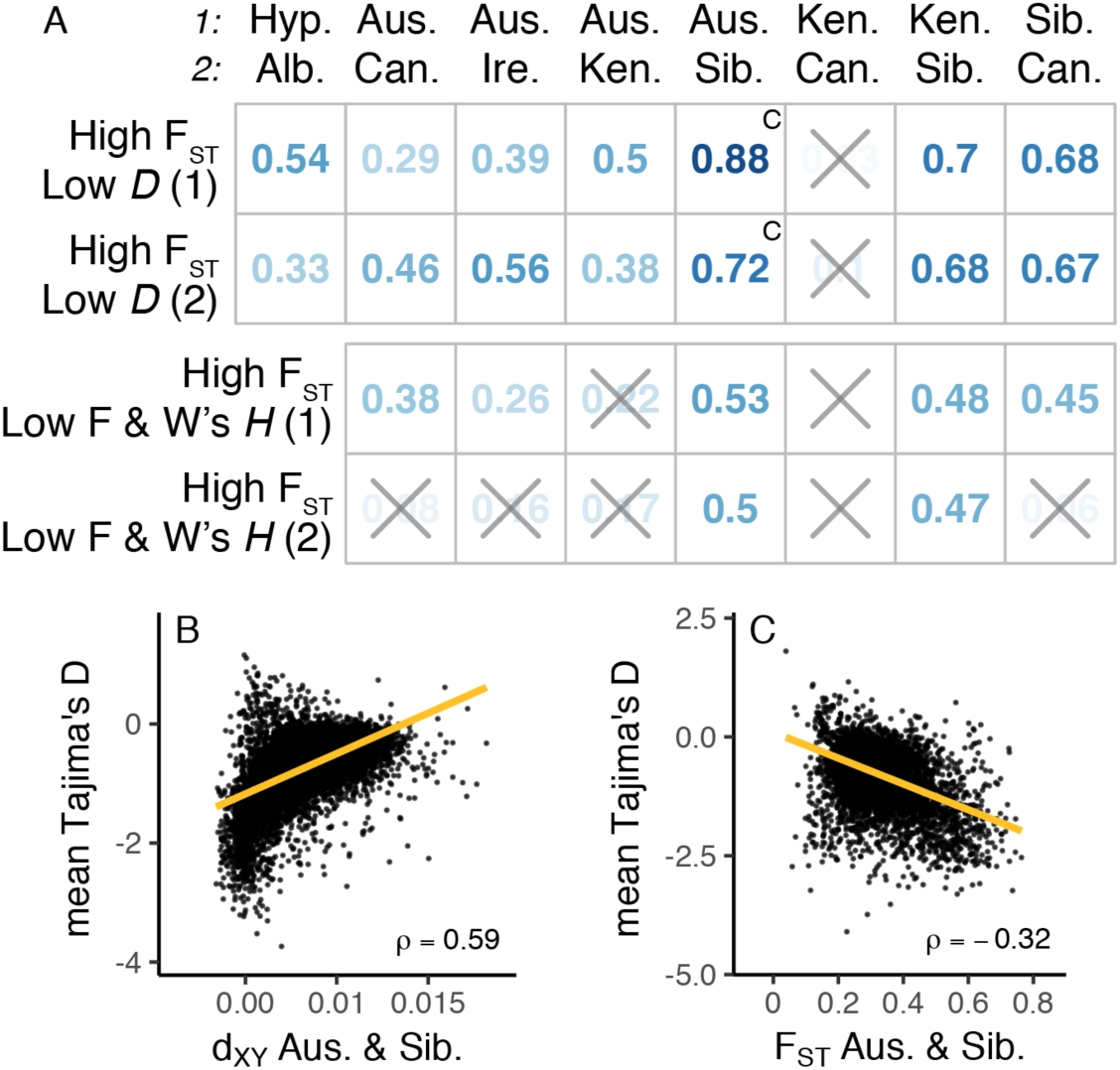
Correlation of F_ST_ and d_XY_ with Tajima’s D and Fay & Wu’s H among stonechats and flycatchers (Ficedula hypoleuca and albicollis, or “Hyp.” and “Alb.”). (A) shows outlier similarity scores, which quantify the number of high-F_ST_ “peaks” that coincide with low Tajima’s D (top section) and low Fay & Wu’s H (bottom section). Within each section, the top (No. 1) and bottom (No. 2) rows show the results for each of the two taxa being compared. This is necessary because Tajima’s D and Fay & Wu’s H are single-population statistics, while ^F^ST and d_XY_ compare two populations. (B,C) show scatterplots where each point represents one 50-Kb genomic window. Refer to Figs. 2–3 for details.

The genomic baseline value of Tajima’s *D* can be biased downward by demographic effects, particularly a population expansion. All stonechat taxa had median Tajima’s *D* between −0.5 and −1.1, with the exception of Canary Islands stonechats, at −2.8 (Fig. 9). Negative values suggest that all five stonechat taxa have experienced past demographic expansion events, with the signal especially strong in the insular Canary Islands stonechats.

**Figure 9.**
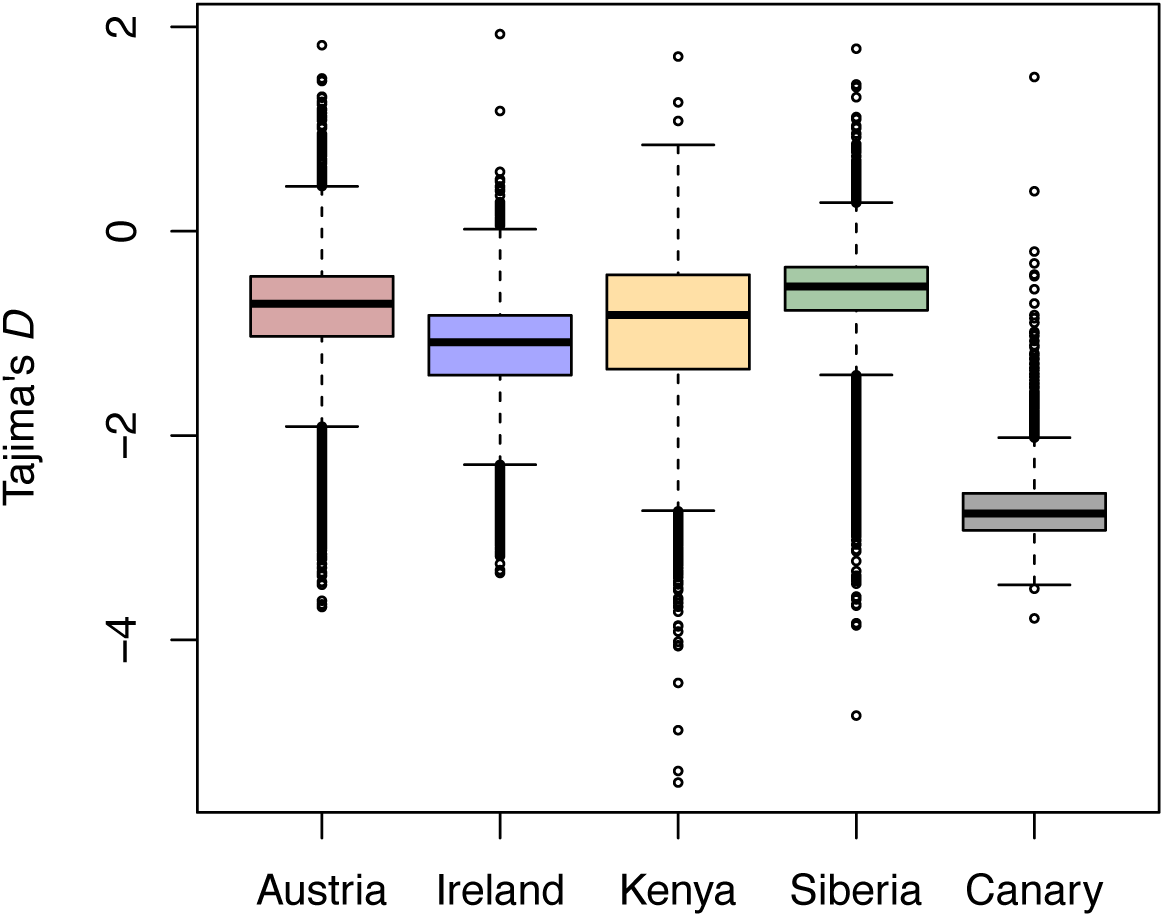
Boxplot of Tajima’s D for five stonechat taxa. Each data point represents one 50-Kb window. Canary Islands stonechats showed the lowest median Tajima’s D.

### Correspondence of genomic landscapes between *Saxicola* and *Ficedula*

Genome-wide patterns of genetic diversity and differentiation were also correlated between stonechats and flycatchers. Absolute divergence was correlated between the two genera (ρ = 0.37-0.49, Fig. 2 C-D); d_XY_ outlier similarity was 0.48-0.52 between stonechats and flycatchers. Flycatchers and stonechats shared a significant number of F_ST_ peaks and valleys, but only for a subset of stonechat comparisons (Fig. 3 E). Genome-wide correlations of F_ST_ were significant but weak (Fig. 3 F-G), and the strongest correlations occurred with Siberian stonechats. Finally, within-population genetic diversity (π) was strongly correlated between stonechat and flycatcher populations; some stonechat-flycatcher correlations were as strong or stronger than stonechat-stonechat correlations (Fig. 6 E-G). Overall, these results suggest that common processes are working independently and in parallel to influence genetic variation in similar regions of the genome in both genera, although the association between genera is weaker than within *Saxicola*.

## DISCUSSION

We examined patterns of genetic diversity and differentiation in an avian radiation and found evidence of selection in similar genomic regions of independent lineages. We identified regions characterized by low within-population diversity, low absolute inter-taxon divergence, and high (or, in some cases, low) differentiation. We found that many stonechat outlier regions also appeared in the closely related genus *Ficedula*. In this genus, genomic regions of low genetic diversity and high differentiation are associated with infrequent recombination (Burri et al. 2015), which suggests that one possible explanation for the parallel patterns of differentiation in these genera is conserved (or convergently evolving) variation in recombination rate (see Singhal et al. 2015). Overall, our results are consistent with linked selection (positive selective sweeps and/or background selection) shaping large-scale patterns of genomic variation in Muscicapid birds. The presence of Fay and Wu’s *H* valleys in differentiation outlier regions supports a role of positive selection in at least some cases. Despite a strong signal of similarity in genomic landscapes, we also found evidence for substantial lineage-specific evolution: Siberian stonechats appear to have experienced the strongest effects of selection, while drift may have shaped Canary Islands stonechats’ genomes.

### Phylogeny

The phylogenetic tree constructed with SNPs from across the nuclear genome (Fig. 1A) was highly supported at all nodes, yet it is not fully concordant with previous trees constructed from mitochondrial DNA sequences (Illera et al. 2008; Woog et al. 2008; Zink et al. 2009). These placed Kenyan stonechats (instead of Siberian, as in our reconstruction) as the sister lineage to the remaining stonechats. Branch support for a sister relationship of Siberian stonechats and the European/Canary Islands clade varied by study and tree-building algorithm. Mito-nuclear discordance could be a sign of past admixture, sex biased gene flow, or other biological phenomena (see Toews and Brelsford 2012). This well-resolved nuclear phylogeny serves as a basis for testing broader questions about genome-scale differentiation in this complex: For example, it helps explain why mean d_XY_ between European and Kenyan stonechats is relatively low compared to Siberian stonechats (Fig. 1C). Finally, although sparse taxon sampling (5 taxa) and choice of outgroup could potentially introduce biases (e.g., Stervander et al. 2015), our cytochrome *b*-only tree (not shown) was consistent with previous mitochondrial studies, which achieved near-complete taxon sampling (e.g., Illera et al. 2008). The topology differences we find between mitochondrial and nuclear-based phylogenies are therefore unlikely to be artifacts of sampling.

### Congruent genomic landscapes across a speciation continuum

Patterns of within- and between-population genetic diversity in stonechats show high levels of parallelism across multiple scales of evolutionary divergence. We found outliers in comparisons of highly similar taxa in the same regions as comparisons at deeper levels of divergence. The parallel reductions in d_XY_ are highly suggestive of selection before divergence (Cruickshank and Hahn 2014), and analogous patterns in standardized nucleotide diversity (π/d_XY_) indicate that common selective forces have continued to reduce diversity on the branches leading to present-day taxa (see Irwin et al. 2016).

Reductions in Fay and Wu’s *H* in some outlier regions suggest that positive selection has played a role in driving some of these regions of low genetic diversity and high differentiation. Some *H* outliers are present in multiple taxa, while others occur in only one (as in *Ficedula*, Burri et al. 2015), suggesting that localized selective sweeps may not have occurred in all groups.

Pairwise comparisons that include Siberian stonechats show the most conspicuous F_ST_ peaks, which coincide with regions of low within-population diversity (π). Together, strongly reduced within-population genetic diversity in specific genomic regions and corresponding peaks of relative differentiation are consistent with Siberian stonechats experiencing the strongest effects of selection in outlier regions. Most of the larger outlier regions also showed significant decreases in Fay and Wu’s *H*, suggesting that positive selective sweeps have contributed to this pattern.

Kenyan and Canarian stonechats showed the highest genome-wide F_ST_. Notably, however, we also observed conspicuous F_ST_ *valleys* in the same locations as the F_ST_ peaks of other pairwise taxon comparisons (Fig. 7 and Fig. S5, Supporting information). In other words, these taxa are differentiated across the vast majority of the genome, but they show low differentiation (F_ST_) in regions of low absolute divergence (d_XY_). This pattern is not unique to this comparison: Austrian and Irish stonechats, and occasionally others, show valleys in similar areas. F_ST_ valleys may occur where d_XY_ (between-group variation) is reduced but π (within-group variation) remains high, especially in Canary Islands stonechats, but more work is needed to understand this phenomenon.

### Correlation of genomic variation between genera

Genomic landscapes of genetic diversity and differentiation in stonechats are significantly correlated with those in Pied and Collared Flycatchers. These results contrast with recent findings in other passerine birds, for example greenish warblers (Irwin et al. 2016). Nucleotide diversity in greenish warblers is only weakly correlated with that in closely related outgroup comparisons (π: Pearson’s r = 0.19). We found a stronger association in nucleotide diversity between stonechats and flycatchers (π: Spearman’s ρ = 0.47-0.60, excluding Canary Is.). *Saxicola* and *Ficedula* share certain aspects of their life history (e.g., they are insectivores, and the flycatchers and most stonechats are migratory), but the hypothesis that these parallel signatures of selection and differentiation are due to shared ecological selection pressures on the same loci appears unlikely. Burri et al. (2015) demonstrated a clear link between low recombination and areas of high divergence in *Ficedula*, which suggests that low recombination might also contribute to shared differentiation outliers within *Saxicola.* Although initial evidence suggested that avian recombination landscapes change drastically over time (Backström et al. 2010), recent work has shown that recombination landscapes can be conserved in birds across millions of years of evolution (Singhal et al. 2015). It is therefore possible that coincident areas of low recombination, in combination with linked selection, may play a role in shaping the broad patterns of landscapes of genomic variation and differentiation across both closely related and deeply diverged taxa. However, direct measures of recombination rates in stonechats are needed to test this hypothesis. While recombination is reduced in close proximity to avian centromeres (Backström et al. 2010), centromeres do not explain the recombination deserts in the centers of acrocentric chromosomes (e.g. 4A, 9, 10, 11, 12, 13, and 18 (Knief and Forstmeier 2015)) (Kawakami et al. 2014; Burri et al. 2015). These regions frequently show high differentiation among flycatchers (Burri et al. 2015) and stonechats.

Decreases in Fay and Wu’s *H* in a subset of outlier regions and a subset of taxa suggest that positive selection has also contributed to this convergent genomic evolution. Indeed, Irwin et al. (2016) favor positive selection as the likely driver of divergence landscapes in greenish warblers, citing exceedingly low nucleotide diversity in differentiation peaks; in one comparison in that study, regions with F_ST_ > 0.9 showed just 6.7% the nucleotide diversity of regions with F_ST_ < 0.6. We found diversity reductions in stonechats and flycatchers to be less severe: between Austrian and Siberian stonechats, which show the greatest reduction in nucleotide diversity in F_ST_ peaks, π in regions with F_ST_ above the 95% percentile was reduced to 30-34% of that of regions with F_ST_ below the 50^th^ percentile. In *Ficedula* flycatchers, this statistic was 43-50%. Therefore, we consider background selection, in concert with reduced recombination, to be an additional plausible driver of correlation in genomic landscapes.

Conserved variation in mutation rate is another possible driver of this correlation. Irwin et al. (2016) found weak correlations in d_XY_ between greenish warblers and more distant comparisons (Pearson’s r = 0.07-0.14), which does not support this explanation. In contrast, we found reasonably strong correlations in d_XY_ between stonechat and flycatcher genera (Spearman’s ρ = 0.37-0.49). Therefore, we cannot rule out a further contribution of conserved variation in mutation rate to these patterns.

### Genomics and demography of Canary Islands stonechats

Canary Islands stonechats’ genomic landscapes differed from those of the other stonechats. The valleys of standardized nucleotide diversity (π/d_XY_) seen in other taxa were completely absent, suggesting that selection has not reduced diversity across the genome in a detectable way. Tajima’s *D* was highly negative genome-wide and showed lower variance than in other stonechats. Combined, these results are most consistent with a demographic history that included a severe population bottleneck (erasing existing patterns of variation), followed by a substantial population expansion. Previous research has found evidence of founder effects and/or bottlenecks in Canary Island birds (Barrientos et al. 2009; Barrientos et al. 2014; Spurgin et al. 2014). The evidence for a bottleneck and expansion and the marked homogeneity of genetic diversity across the genome in Canary Islands stonechats suggest that genetic drift has played a dominant role in its divergence from other stonechats, possibly overpowering selection (see Hansson et al. 2014; Spurgin et al. 2014; Gonzalez-Quevedo et al. 2015; Illera et al. 2016).

### Evidence of lineage-specific evolution

Despite striking similarities in the genomic landscapes of stonechats, we also find lineage-specific evolution. At the broadest levels of our analysis, in which we compare genera, we identified conspicuous divergence peaks that appear in *Ficedula* but not *Saxicola* (e.g., on chromosomes 3, 8, 10, 11, 12, 13, and 18; Fig. S11, Supporting information, shows a close up of chromosome 13), and vice versa (e.g., on chromosomes 6, 7, 17, and 20; Fig. S12, Supporting information shows chromosome 20). These outlier regions should be further examined from a functional perspective, as they appear to have resulted from evolutionary processes specific to a particular lineage. The most conspicuous outlier regions shared between these systems should likewise be examined (e.g., on chromosomes 1, 1A, 2, 3, 4, and 4A; Figs. S13 and S14, Supporting information, show chromosomes 1A and 4A).

### Conclusion

Few former studies (Burri et al. 2015; Lamichhaney et al. 2015; Irwin et al. 2016) have examined genome-wide patterns of divergence in more than two avian taxa, yet comparative studies of closely related species have great potential to shed light on genome evolution (Cutter and Payseur 2013). We find parallel patterns of selection in the stonechat complex—likely occurring both before and after speciation—and evidence of demography potentially overwhelming signatures of selection in one species. In addition, this study suggests that parallel genomic processes are operating in independent evolutionary systems to drive the differentiation of similar genomic regions across genera. We hypothesize that linked selection coupled with areas of low recombination, which may be conserved across these taxa, have shaped these broad patterns. Whether concordant outlier regions actually contribute to reproductive isolation or are otherwise consequential in the speciation process is unknown. Therefore, we recommend that outlier markers obtained through genome scans and their relevance to speciation be interpreted with caution. Importantly, our comparative method also identified divergence outlier regions that are not widely shared; these may harbor loci important in lineage-specific evolution. As genomic comparisons among radiations accumulate, we will be able to compare the congruence in genomic landscapes and potentially reveal the phenomena that drive genomic divergence over evolutionary time.

## ACKNOWLEDGEMENTS

We thank the Max Planck Society, the Fuller Evolutionary Biology program at the Cornell Lab of Ornithology, the Hunter R. Rawlings Presidential Research Scholars Program (Cornell), and the Dextra Undergraduate Research Endowment Fund (Cornell) for funding. We thank the University of Washington Burke Museum, Heiner Flinks and Bea Apfelbeck for providing samples, and the Canarian Government and the Cabildo of Fuerteventura for permission to sample Canary Islands stonechats. We thank Heinke Buhtz, Conny Burghardt, Sven Künzel, Nicole Thomsen, Mayra Zamora Moreno, and Bronwyn Butcher for guidance in the lab; David Toews, Scott Taylor, Jacob Berv, and Kira Delmore for valuable discussion; Kelly Zamudio, Jennifer Walsh, Kristen Ruegg, Sonya Clegg, Eric Anderson, and three anonymous reviewers for thoughtful feedback on this manuscript; and Susan and Daniel Van Doren for valued support.

